# Cyclooxygenase production of PGE_2_ promotes phagocyte control of *A. fumigatus* hyphal growth in larval zebrafish

**DOI:** 10.1101/2021.10.19.464929

**Authors:** Savini Thrikawala, Mengyao Niu, Nancy P. Keller, Emily E. Rosowski

## Abstract

Invasive aspergillosis is a common opportunistic infection, causing >50% mortality in infected immunocompromised patients. The specific molecular mechanisms of the innate immune system that prevent pathogenesis of invasive aspergillosis in immunocompetent individuals are not fully understood. Here, we used a zebrafish larva-*Aspergillus* infection model to identify cyclooxygenase (COX) enzyme signaling as one mechanism that promotes host survival. Larvae exposed to the pan-COX inhibitor indomethacin succumb to infection at a significantly higher rate than control larvae. COX signaling is both macrophage- and neutrophil-mediated. However, indomethacin treatment has no effect on phagocyte recruitment. Instead, COX signaling promotes phagocyte-mediated inhibition of germination and invasive hyphal growth. Protective COX-mediated signaling requires the receptor EP2 and exogenous prostaglandin E_2_ (PGE_2_) rescues indomethacin-induced decreased immune control of fungal growth. Collectively, we find that COX signaling activates the PGE_2_-EP2 pathway to increase control *A. fumigatus* hyphal growth by phagocytes in zebrafish larvae.

**Author Summary:** Invasive aspergillosis causes mortality in >50% of infected patients. It is caused by a free-living fungus *Aspergillus fumigatus* which releases thousands of airborne spores. While healthy individuals clear inhaled spores efficiently, in immunocompromised individuals these spores grow into filamentous hyphae and destroy lungs and other tissues causing invasive aspergillosis. The immune mechanisms that control this fungal growth in healthy people are still largely unknown. Here, we used a larval zebrafish model of *A. fumigatus* infection to determine that cyclooxygenase enzymes, which are the target of non-steroidal anti-inflammatory drugs such as aspirin and ibuprofen, are important to control the fungus. Innate immune cells use cyclooxygenase signaling to prevent hyphal growth and tissue destruction. Our study provides new insights into the mechanisms that immune cells deploy to stop invasive growth of *A. fumigatus* and inform development of future strategies to combat invasive aspergillosis.

## Introduction

*Aspergillus fumigatus* is a free-living saprophytic fungus which reproduces asexually by producing thousands of conidia or spores. Owing to their small size and hydrophobicity, spores become airborne causing widespread contamination both indoors and outdoors. It is estimated that an average person can inhale 100-1000 spores per day (1). Although healthy immune systems can combat these spores, in immunocompromised individuals spores can germinate to form invasive hyphae which spread to multiple organs and tissues—a condition called invasive aspergillosis (IA) (1). IA remains a major cause of mortality in immunocompromised patients, particularly individuals with hematological malignancies, bone marrow or solid-organ transplant recipients, HIV patients, ICU patients, and patients with altered lung conditions (2). Despite the availability of anti-fungal drugs, the mortality rate of IA remains at ~50% (3–5). Hence, it is imperative to develop novel strategies to target fungi and augment anti-fungal immune responses, but this requires a better understanding of immune cell-pathogen interactions. Innate immune cells act as the first line of defense against inhaled *A. fumigatus* conidia. However, the signaling and effector mechanisms that these cells use to inhibit fungal growth are not fully understood.

Eicosanoids, such as prostaglandins, are arachidonic acid-derived lipid signaling molecules that function in both an autocrine and paracrine manner by binding to their receptors and can have a variety of effects on immune cell function (6, 7). Prostaglandins are produced by prostaglandin endoperoxide H synthases (PGHSs), also called cyclooxygenase (COX) enzymes. COX enzymes are the target of non-steroidal anti-inflammatory drugs such as aspirin, ibuprofen and indomethacin (8–10). COX-derived prostaglandins can have either pro- or anti-inflammatory effects on innate immune cells, modulating both phagocyte recruitment and phagocyte functions (6, 7, 11). In response to fungal pathogens, prostaglandin signaling is known to inhibit phagocytosis of *Candida albicans* by macrophages, H_2_O_2_-mediated fungicidal activity against *Paracoccidioides brasiliensis,* and M1 polarization of alveolar macrophages and killing of *Cryptococcus neoformans* (12–14). However, the roles of COX activation and prostaglandins during *A. fumigatus* infections are not known.

Analyzing how given pathways affect specific aspects of dynamic host-pathogen interactions *in vivo* is challenging. The zebrafish larva-*Aspergillus* infection model overcomes many of these challenges, as larvae are transparent and allow for direct visualization of phagocyte-*Aspergillus* interactions through high-resolution repetitive imaging of the same larvae over the course of a multi-day infection (15). Multiple steps in pathogenesis such as phagocyte recruitment, phagocytosis, spore killing, germination, and hyphal growth or clearance can be quantified using this live imaging technique (16). Zebrafish have a well-conserved immune system with humans, but depend solely on their innate immune system for the first few weeks of their life, providing a window to study innate immune mechanisms with no interference from the adaptive system (17, 18). The zebrafish larva-*Aspergillus* model recapitulates multiple aspects of human IA: while immunocompetent larvae are resistant, immunocompromised larvae are susceptible to the infection and develop invasive hyphae (19, 20).

Here we use this zebrafish larva-*Aspergillus* infection model to determine the role of the host COX pathway in phagocyte-mediated *A. fumigatus* clearance. We find inhibition of host COX signaling increases spore germination and invasive growth of hyphae in infected larvae, thereby decreasing host survival. COX signaling does not affect macrophage or neutrophil recruitment but instead activates these cells to target the fungus. Exogenous PGE_2_ injection restores control of hyphae in COX-inhibited larvae, suggesting that PGE_2_ is a major driver of COX-mediated control of fungal growth by phagocytes.

## Results

### Host cyclooxygenase inhibition decreases infected larval survival

Prostaglandins are lipid signaling molecules whose production is induced during inflammation via cyclooxygenase (COX) enzymes. We used the zebrafish larva-*Aspergillus* infection model to test the hypothesis that host COX signaling promotes larval survival and fungal clearance in an *A. fumigatus* infection. Wild-type *A. fumigatus* spores were microinjected into the hindbrain ventricle of 2 days post fertilization (dpf) larvae. Infected larvae were then exposed to the pan-COX inhibitor indomethacin or DMSO vehicle control immediately after injection and larval survival was monitored for 7 days. Indomethacin is a well-established non-steroidal anti-inflammatory drug that inhibits COX enzyme activation and prostaglandin synthesis (9, 10) and is widely used in a variety of animal models including zebrafish (21, 22). Indomethacin-treated larvae succumb to infection at a significantly greater rate than control larvae (Fig 1A). Treatment with the COX1 inhibitor SC560 (23) or COX2 inhibitor meloxicam (21) also significantly increases infected larval mortality (S1 Fig). With each of these inhibitors, no significant decrease in survival was observed in mock-infected larvae (Fig 1A and S1 Fig).

**Fig. 1.**
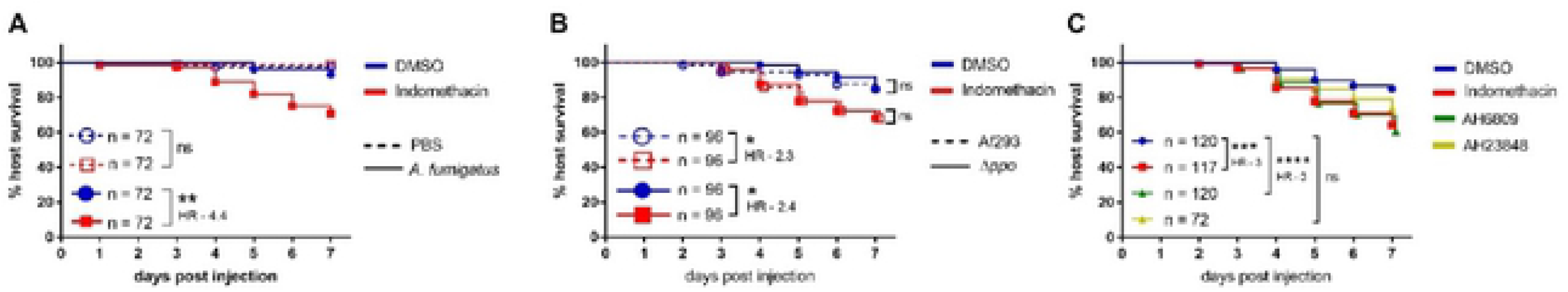
Host cyclooxygenase signaling promotes survival of *A. fumigatus*-infected larvae. **(A)** Survival of wild-type larvae injected at 2 dpf with TBK1.1 (Af293) *A. fumigatus* spores or PBS mock injection in the presence of 10 μM indomethacin or DMSO vehicle control. **(B)** Survival of larvae injected at 2 dpf with *A. fumigatus* Af293 or Af293 triple-*ppo*-mutant (Δ*ppo*) spores and exposed to 10 μM indomethacin or DMSO vehicle control. **(C)** Survival of larvae injected at 2 dpf with TBK1.1 (Af293) spores and exposed to 10 μM indomethacin, 5 μM AH6809, 10 μM AH23848, or DMSO vehicle control. Data are pooled from at least three independent replicates and the total larval N per condition is indicated in each figure. Cox proportional hazard regression analysis was used to calculate P values and hazard ratios (HR). Average injection CFUs: (A) 50, (B) Af293 = 40 and Δ*ppo* = 58, (C) 28.

*A. fumigatus* also harbors three Ppo enzymes (PpoA, PpoB and PpoC) with high identity to vertebrate COX (24), but a previous study reported that indomethacin does not affect the function of these enzymes (25). To confirm that the observed effects of indomethacin on infected larval survival are not due to inhibition of fungal enzymes, we infected zebrafish larvae with *A. fumigatus* spores lacking all three *ppo* genes (Δ*ppoA*, Δ*ppoB*, Δ*ppoC*) (S2 Fig). Deletion of *ppo* enzymes had no effect on fungal virulence, as survival of larvae infected with triple-*ppo*-mutant *A. fumigatus* spores is similar to larvae infected with wild-type spores (Fig 1B). Additionally, indomethacin treatment decreased survival equally in larvae infected with triple-*ppo*-mutant and wild-type spores (Fig 1B). These data demonstrate that indomethacin inhibits host enzymes to compromise survival of *A. fumigatus*-infected larvae. Since no survival difference was observed in larvae infected with a triple-*ppo*-mutant, we focused on wild-type *A. fumigatus* for the remainder of the study.

Among COX-biosynthesized prostaglandins, prostaglandin E_2_ (PGE_2_) is a major product synthesized by phagocytes that moderates a range of inflammatory processes and has pro- or anti-inflammatory functions, depending on the receptor to which it binds (26). PGE_2_ elicits its actions via four different E type prostanoid receptors, EP1-4, with most immunomodulatory effects mediated via EP2 and EP4 (26). Therefore, we tested if EP2 and 4 receptor antagonists affect the disease outcome of *A. fumigatus*-infected larvae. We used antagonists of EP2: AH6809 (27) and EP4: AH23848 (22, 27, 28) previously used in zebrafish larvae. Larvae exposed to AH6809 succumb to the infection at a similar rate as indomethacin-exposed larvae, both with a hazard ratio of 3 compared to control larvae, while AH23848-exposed larvae show no significant difference in survival compared to control (Fig 1C). These data suggest that COX signaling promotes *A. fumigatus*-infected larvae survival via a PGE_2_-EP2 signaling pathway.

### Both macrophages and neutrophils use cyclooxygenase signaling to combat *A. fumigatus* infection

We next sought to determine which innate immune cells utilize COX signaling to fight *A. fumigatus* infection. Macrophages and neutrophils are the primary immune cells that combat *A. fumigatus* infection in zebrafish larvae (20). To determine if these cell types play a role in COX-mediated host responses, we inhibited development of both phagocytes by knocking down *pu.1 (spi1b)* via morpholino injection (29). If COX signaling activates phagocytes to clear the infection, we expect that indomethacin treatment of larvae that are already depleted of phagocytes would have no effect on larval survival. Larvae were injected with *A. fumigatus* spores or PBS and exposed to indomethacin or DMSO. Indomethacin exposure significantly decreases survival of larvae injected with a control morpholino but has no effect on *pu.1* morphants (Fig 2A, S3A Fig), suggesting that COX-mediated host protection is phagocyte-dependent.

**Fig. 2.**
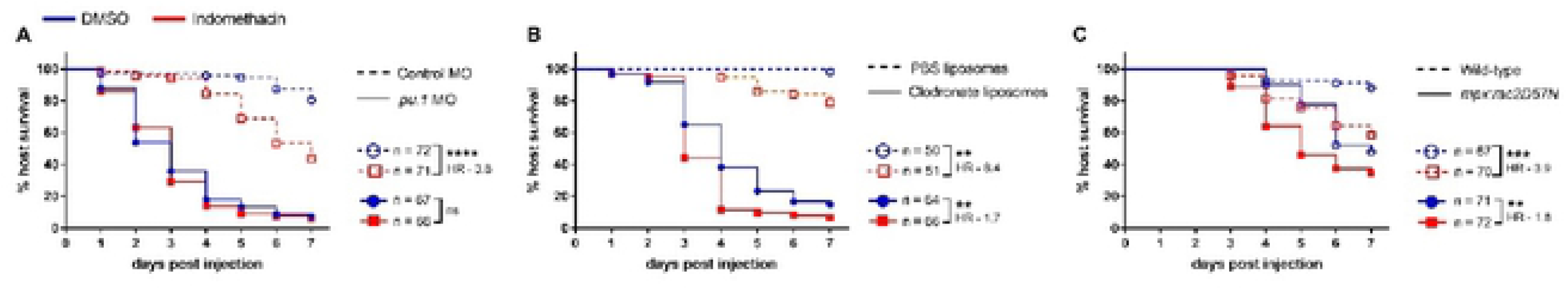
Cyclooxygenase-mediated host protection depends on phagocytes. Survival of larvae injected with TBK1.1 (Af293) spores at 2 dpf. **(A)** Development of phagocytes was inhibited by *pu.1* morpholino (MO). Control larvae received standard control MO. **(B)** Macrophages were depleted via clodronate liposome i.v. injection at 1 dpf. Control larvae received PBS liposomes. **(C)** Neutrophil-defective larvae (*mpx*:*rac2D57N*) were compared to wild-type larvae. Data are pooled from three independent replicates and the total larval N per condition is indicated in each figure. Cox proportional hazard regression analysis was used to calculate P values and hazard ratios (HR). Average injection CFUs: (A) control MO = 25, *pu.1* MO = 24, (B) PBS liposomes = 20, clodronate liposomes = 23, (C) wild-type = 26, *mpx*:*rac2D57N* = 22.

Then we interfered with macrophage and neutrophil function individually to determine if each cell type is required for COX-mediated host protection. We injected 1 dpf larvae with clodronate liposomes to deplete macrophages or PBS liposomes as a control. As observed previously, macrophage-depleted larvae rapidly succumb to the infection (20). However, indomethacin exposure further decreases the survival of macrophage-depleted larvae (Fig 2B). While indomethacin treatment makes control larvae 8.4 times more likely to succumb to infection, clodronate liposome-injected larvae are only 1.7 times more likely to succumb upon indomethacin treatment (Fig 2B), suggesting that macrophages partially mediate the host-protective effects of COX signaling, but that even in the absence of macrophages COX signaling increases host survival. Larvae injected with clodronate liposomes and then given a PBS mock infection also have lower survival upon indomethacin treatment, suggesting that some of this death may be due to the effects of the clodronate alone, although this difference in PBS mock-infected larvae is not statistically significant (S3B Fig).

We next tested the survival of neutrophil-defective (*mpx:rac2D57N*) infected larvae. In these larvae neutrophils are unable to migrate to the infection site (30). As found previously, neutrophil-defective larvae are more susceptible to *A. fumigatus* infection than wild-type controls (Fig 2C) (20, 31). Indomethacin exposure further decreases survival of neutrophil-defective larvae (Fig 2C). Compared to wild-type larvae which are 3.9 times more likely to succumb to infection, neutrophil-defective larvae are only 1.8 times more likely to succumb to infection, suggesting that neutrophils also partially mediate the host-protective effects of COX signaling, but that other cell types can be involved. Lack of neutrophils has no effect on survival of mock-infected larvae treated with indomethacin (S3C Fig). Together, these data demonstrate that both macrophages and neutrophils participate in COX-mediated responses to promote survival of *A. fumigatus*-infected zebrafish larvae.

### Cyclooxygenase activity is not required for phagocyte recruitment

We next sought to define how the innate immune response is altered by COX inhibition. COX-synthesized prostaglandins are chemical messengers that can function to recruit immune cells to infection sites, and we wondered whether COX inhibition affects macrophage or neutrophil recruitment to *A. fumigatus* infection (7, 20). Zebrafish larvae expressing GFP in macrophages (*Tg(mpeg1:H2B-GFP)*) or BFP in neutrophils (*Tg(lyz:BFP)*) were infected with *A. fumigatus* spores expressing mCherry, and treated with indomethacin or DMSO vehicle control and we enumerated the number of macrophages and neutrophils at the infection site through daily confocal imaging (Fig 3A).

**Fig. 3.**
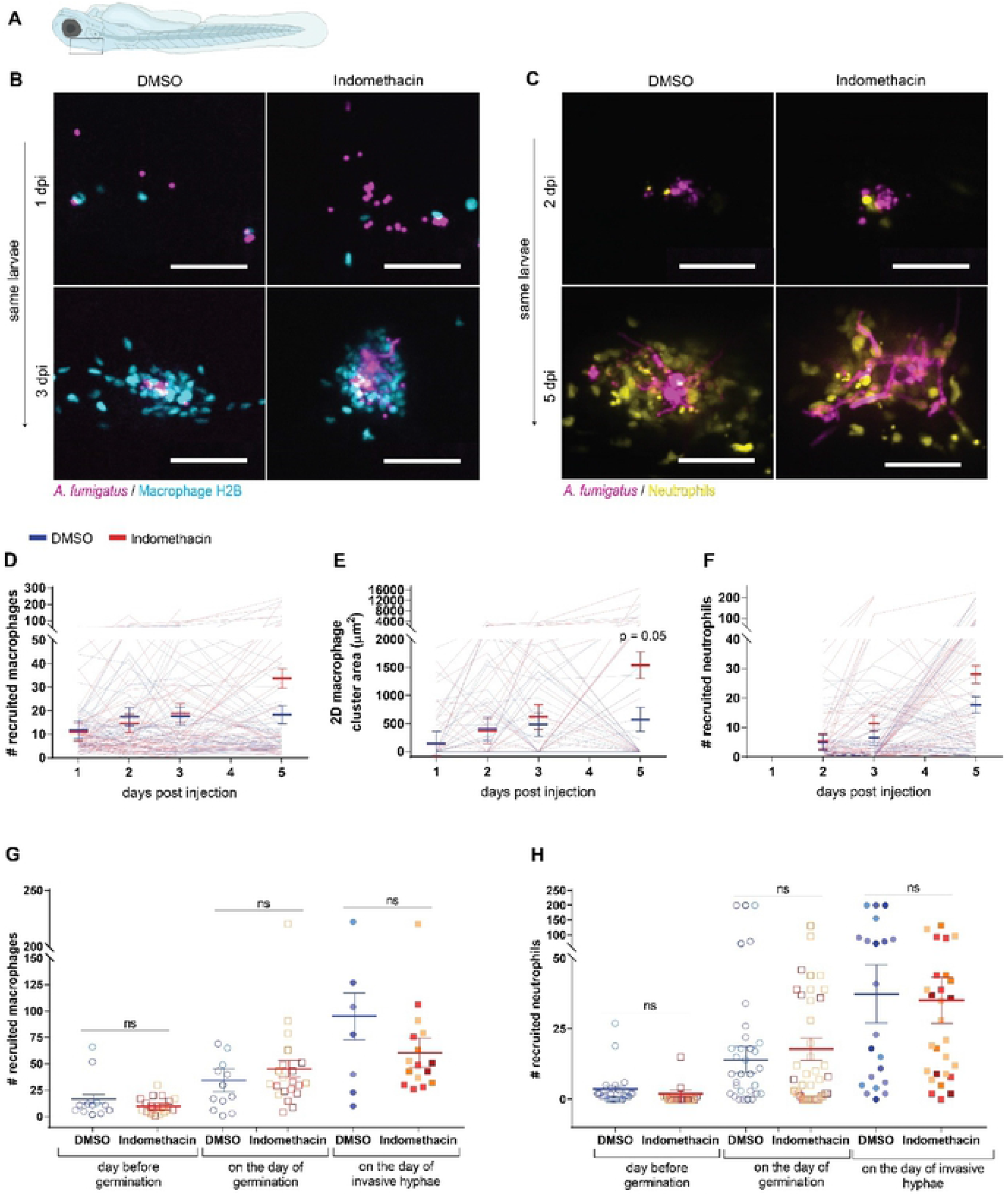
Cyclooxygenase inhibition does not alter phagocyte recruitment. Larvae were injected with mCherry-expressing *A. fumigatus* TBK5.1 (Af293) spores at 2 dpf. After injection larvae were exposed to 10 μM indomethacin or DMSO vehicle control and were imaged through 5 dpi. **(A)** Schematic showing the infection and imaging area of zebrafish larvae. **(B, D, E, G)** Macrophage nuclear-labeled *Tg(mpeg1:H2B-GFP)* larvae were imaged at 1, 2, 3, and 5 dpi. **(C, F, H)** Neutrophil-labeled *Tg(lyz:BFP)* larvae were imaged at 2, 3, and 5 dpi. (B, C) Representative z-projection images showing macrophage and neutrophil recruitment. Scale bars = 50 μm. (D) Number of macrophages recruited, (E) 2D macrophage cluster area, and (F) number of neutrophils recruited were quantified. Each line represents an individual larva followed for the entire course of infection and bars represent pooled emmeans ± SEM from four independent replicates, at least 12 larvae per condition, per replicate. P values were calculated by ANOVA. (G) Number of macrophages and (H) neutrophils one day before germination occurred, on the day of germination, and on the day invasive hyphae occurred were quantified. Bars represent pooled emmeans ± SEM from all larvae with germination from four independent replicates. Data points represent individual larva and are color coded by replicate. P values were calculated by ANOVA.

As described previously, macrophages are recruited starting at 1 day post infection (dpi) and form clusters around spores starting at 2-3 dpi (Fig 3B), with neutrophils primarily responding after spores germinate (Fig 3C). The number of recruited macrophages (Fig 3D), macrophage cluster area (Fig 3E), and the number of recruited neutrophils (Fig 3F) are not significantly different between indomethacin- and DMSO-exposed groups at days 1, 2, and 3 post infection. At 5 dpi, however, more macrophages (Fig 3D) and neutrophils (Fig 3F) are found at the infection site in indomethacin-treated larvae. Fungal germination occurs at these later time stages and attracts more immune cells. To control for this variable, we analyzed the number of macrophages and neutrophils in each larva relative to the day germination and invasive hyphae were first observed. Using this normalization, we find that macrophage numbers (Fig 3G) and neutrophil numbers (Fig 3H) are similar between the two conditions at each stage of fungal pathogenesis. Overall, our results indicate that phagocyte recruitment is not dependent on COX activation.

### Cyclooxygenase activity does not promote spore killing

We next hypothesized that the functions of these phagocytes are modulated by COX signaling. The initial response of macrophages is to phagocytose injected spores and activate spore killing mechanisms (19, 20). To determine if COX inhibition affects spore killing, we used a live-dead staining method in which *A. fumigatus* spores expressing YFP are coated with AlexaFluor 546 and injected into zebrafish larvae expressing mTurquoise in macrophages (20, 32). Larvae were imaged with confocal microscopy at 2 dpi, and we enumerated the number of live versus dead spores. Live spores are visualized as YFP signal surrounded by AlexaFluor signal, while dead spores only have AlexaFluor signal (Fig 4A). The percentage of live spores is similar in indomethacin and DMSO groups both within macrophages and in the whole imaged hindbrain area (Fig 4B). To confirm these results, we also measured the overall fungal burden in indomethacin- or DMSO-treated larvae over the 7-day infection period with CFU counts. Consistent with live-dead staining, the fungal burden is similar between DMSO- and indomethacin-exposed larvae throughout the infection (Fig 4C), indicating that COX signaling does not drive spore clearance.

**Fig. 4.**
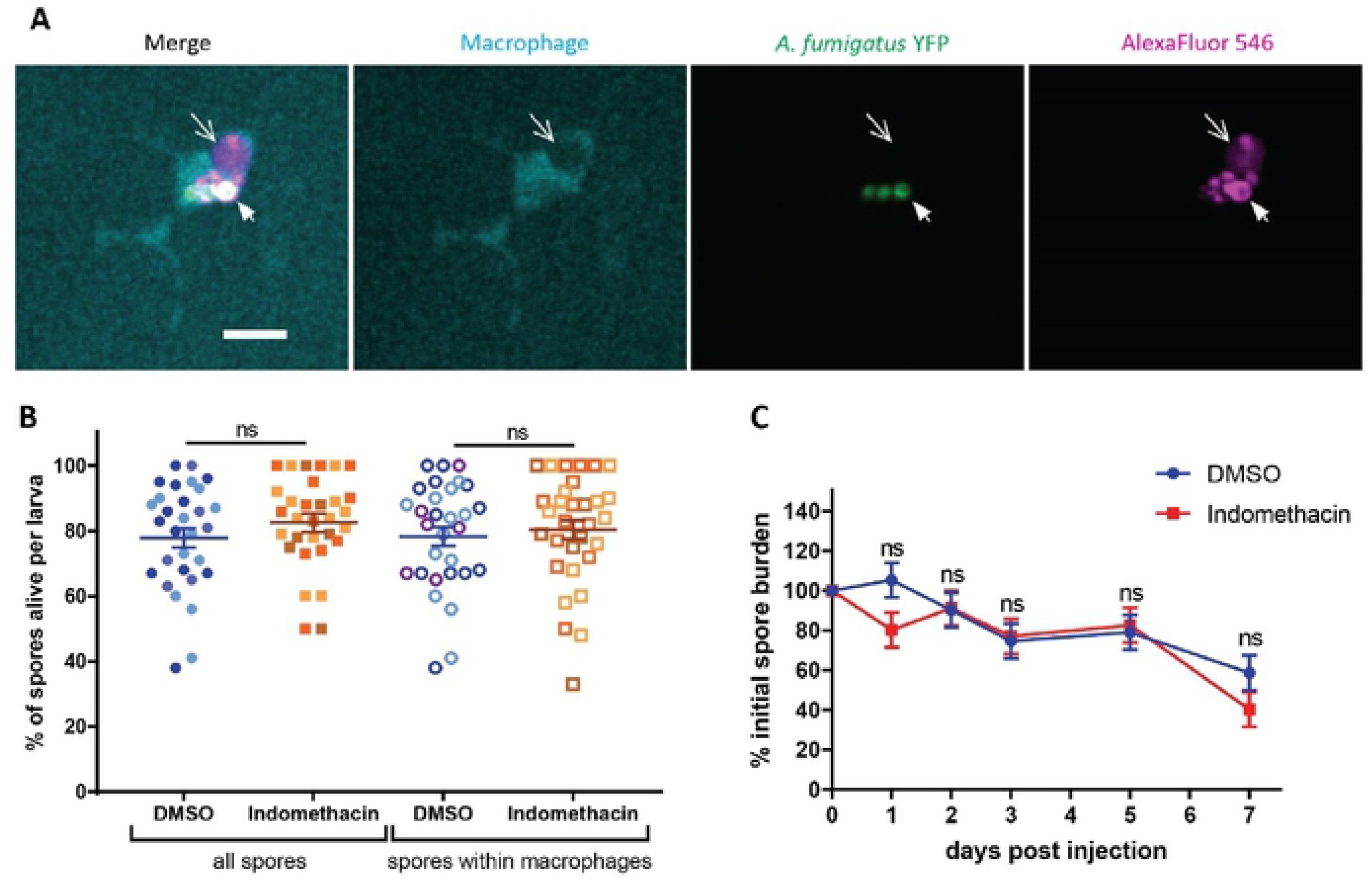
Cyclooxygenase inhibition does not affect spore killing. **(A, B)** Macrophage-labeled larvae *Tg(mfap4:mTurquoise2)* were injected with YFP-expressing *A. fumigatus* TBK1.1 (Af293) spores coated with AlexaFluor 546 at 2 dpf, exposed to 10 μM indomethacin or DMSO vehicle control, and imaged at 2 dpi. (A) Representative images showing live (white arrow) and dead (open arrow) spores within a macrophage. Scale bar = 10 μm. Z projection of three slices. (A) The percentage of live spores in the hindbrain, and specifically within macrophages, per larvae. Each data point represents an individual larvae, color-coded by replicate (indomethacin n = 31, DMSO n = 30). Bars represent pooled emmeans ± SEM from three independent replicates, P values calculated by ANOVA. **(C)** Wild-type larvae were injected with TBK1.1 (Af293) spores at 2 dpf, exposed to 10 μM indomethacin or DMSO vehicle control, and fungal burden was quantified by homogenizing and plating individual larvae for CFUs at multiple days post injection. Eight larvae per condition, per dpi, per replicate were quantified, and the number of CFUs at each dpi is represented as a percentage of the initial spore burden. Bars represent pooled emmeans ± SEM from three individual replicates, P values calculated by ANOVA. Average injection CFUs: 27.

### Cyclooxygenase inhibition decreases immune control of fungal germination

As spore killing is not affected by COX inhibition, we next hypothesized that immune control of the next stages in fungal pathogenesis—spore germination and invasive hyphal growth—is modulated by COX signaling. To monitor spore germination and hyphal growth in larvae, we infected larvae with *A. fumigatus* spores expressing mCherry and imaged at 1, 2, 3 and 5 dpi with confocal microscopy. We find spore germination in both indomethacin- and DMSO-exposed larvae (Fig 5A). However, both the rate at which larvae are observed to have germination inside of them and the total percentage of larvae that harbor germinated spores is significantly higher in the presence of indomethacin (Fig 5B).

**Fig. 5.**
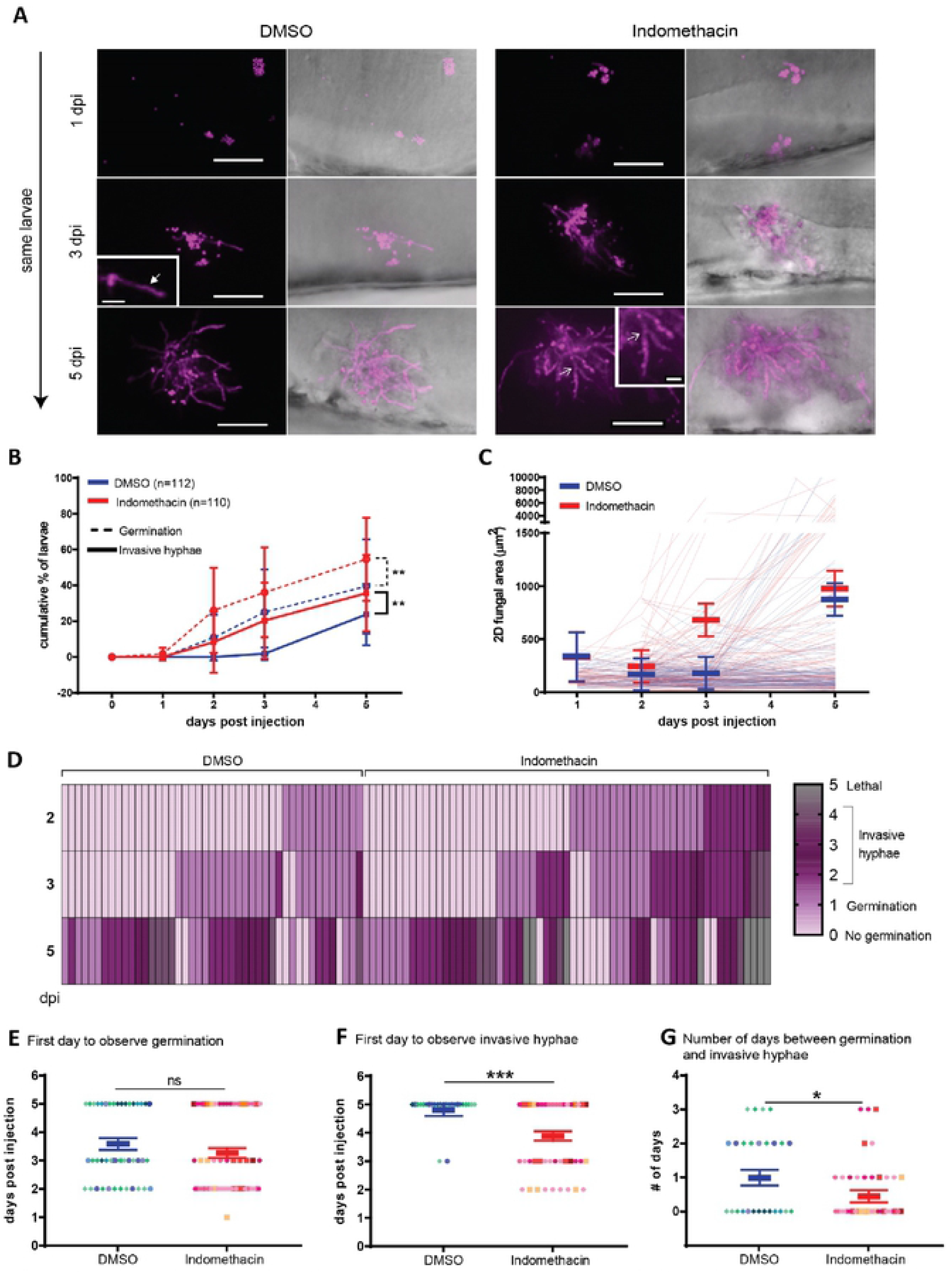
Cyclooxygenase inhibition decreases immune control of fungal germination and invasive hyphal growth. Zebrafish larvae were injected with mCherry-expressing TBK5.1 (Af293) spores at 2 dpf, exposed to 10 μM indomethacin or DMSO vehicle control and imaged at 1, 2, 3, and 5 dpi. **(A)** Representative images showing spore germination (inset white arrow) and invasive hyphae (branched hyphae, inset open white arrow). Scale bar = 50 μm (10 μm in insets). **(B)** Cumulative percentage of larvae with germination (dotted line) and invasive hyphae (solid line) through 5 dpi. Cox proportional hazard regression analysis was used to calculate P values. **(C)** 2D fungal area was quantified from image z projections. Each line represents an individual larva and bars represent pooled emmeans ± SEM from 8 independent replicates, at least 12 larvae per condition, per replicate. **(D)** Severity of fungal growth was scored for all larvae and displayed as a heatmap. Representative images for each score can be found in S5 Fig. **(E-G)** In larvae in which (E) germination (indomethacin n = 61, DMSO n = 45) and (F) invasive hyphae occurred (indomethacin n = 42, DMSO n = 27), the day on which each was first observed is plotted. (G) The number of days between germination and invasive hyphae was also calculated. Bars represent pooled emmeans ± SEM from eight individual replicates, P values calculated by ANOVA. Each data point represents an individual larvae, color-coded by replicate.

Since germination is increased upon indomethacin treatment, we wanted to confirm again that the effects of indomethacin are on the host and that indomethacin does not directly alter *A. fumigatus* germination. To test this, *A. fumigatus* spores were inoculated *in vitro* in liquid RPMI medium in the presence of indomethacin or DMSO and the percentage of germinated spores was scored at 2-hour intervals. We find no difference in the rate of germination between indomethacin and DMSO-treated spores (S4 Fig). Therefore, our data demonstrate that indomethacin decreases host immune cell-mediated control of *A. fumigatus* germination.

### Cyclooxygenase inhibition decreases immune control of invasive hyphal growth

After germination, *A. fumigatus* hyphae branch and grow into a network disrupting the host tissue. The cumulative percentage of larvae with invasive hyphae (as defined by branched hyphal growth (S5 Fig)) is also significantly higher with indomethacin treatment (Fig 5B). We also quantified the hyphal burden by measuring the fungal area, finding more extensive hyphal growth in indomethacin-treated larvae at both 3 and 5 dpi, although this difference is not statistically significantly (Fig 5C), likely due to high variability between larvae and the large number of indomethacin-treated larvae that succumbed to infection before 5 dpi (Fig 1A). Next, we rated the severity of fungal growth on a scale of 0 to 4, from no germination to severe invasive growth of hyphae, and a lethal score of 5 (S5 Fig). Severe growth of invasive hyphae is prominent in larvae exposed to indomethacin, eventually causing mortality (Fig 5D). Although germination occurred in the vehicle control group, these larvae are able to delay invasive growth compared to indomethacin-treated larvae (Fig 5D), suggesting that the major defect in these larvae is a failure to control hyphal growth post-germination. To quantify this time of delay between appearance of germination and invasive hyphae, we analyzed the timeline of first appearance of germlings and invasive hyphae in larvae more closely, focusing only on larvae that had germination within them at some point in the experiment. We quantified the day germination was first observed, the day invasive hyphae was first observed, and the time between these two occurrences. The day to first observe germination was similar between the two groups (Fig 5E). However, invasive hyphae appear significantly earlier after both initial infection (Fig 5F) and after germination (Fig 5G) in indomethacin-exposed larvae. Once spores are germinated, invasive hyphae appear on average ~1 day later in control larvae, while in indomethacin-treated larvae, this growth only takes an average of ~0.5 days (Fig 5G). Together, these results suggest that COX signaling promotes phagocyte-mediated control of invasive hyphal growth.

Neutrophils are thought to be the major phagocyte that targets and kills invasive hyphae. In *irf8*^−/−^ larvae which lack macrophages and have an abundance of neutrophils, neutrophils destroy fast-germinating strains of *A. fumigatus* such as CEA10 within a few days (20). We therefore decided to use this infection scenario to specifically test the requirement for COX signaling in neutrophil-mediated hyphal killing. We infected *irf8*^−/−^ larvae with CEA10 spores and isolated larvae for CFU enumeration at 0, 1, and 2 dpi. CFUs from *irf8*^−/−^ were normalized to CFUs of *irf8*^+/+^/*irf8*^+/−^ at each dpi for each condition. At 2 dpi in DMSO-treated larvae, ~26% of the fungal burden remains in *irf8*^−/−^ larvae compared to macrophage-sufficient larvae (*irf8*^+/+^ or *irf8*^+/−^), demonstrating the neutrophil-mediated clearance of fungus that occurs in these larvae (S6A Fig). Fungal clearance is slightly alleviated but not significantly different in indomethacin-exposed *irf8*^−/−^ larvae, suggesting that COX signaling may promote but is not required for neutrophil-mediated killing of hyphae (S6A Fig). Similar to infection with Af293-derived strains, CEA10-infected larvae also succumb to the infection at a higher rate in the presence of indomethacin both in *irf8*^+/+^/*irf8*^+/−^ and *irf8*^−/−^ backgrounds (S6B Fig).

### Exogenous PGE_2_ increases immune control of hyphal growth in the presence of indomethacin

So far we have established that COX signaling promotes macrophage- and neutrophil-mediated control of germination and invasive hyphal growth in an *A. fumigatus* infection. It is likely that COX signaling acts via a PGE_2_-EP2 signaling axis, as EP2 receptor antagonist-treated infected larvae succumb to infection at the same rate as indomethacin-treated larvae (Fig. 1C). We therefore tested if exogenous PGE_2_ can rescue the effects of indomethacin treatment in infected larvae. Since PGE_2_ is short-lived and elicits short-range effects, we injected *A. fumigatus*-infected, indomethacin- or DMSO-treated larvae with PGE_2_ or DMSO vehicle control into the hindbrain at 1 dpi. PGE_2_ injection partially rescues survival of indomethacin-treated larvae, although the effect is not statistically significant (Fig 6A). To determine if PGE_2_ can rescue indomethacin-inhibited functions of phagocytes against invasive fungal growth, we imaged the larvae at 3 dpi. As seen previously, indomethacin treatment increases the percentage of larvae harboring both germination and invasive hyphae at 3 dpi (Fig 6B). Quantification of 2D fungal area further supports this observation (Figs 6C, D). PGE_2_ supplementation rescues these phenotypes, decreasing germination, development of invasive hyphae, and total fungal burden (Figs 6B-D), without affecting phagocyte recruitment (S7A and S7B Figs). However, in the absence of indomethacin treatment, exogenous PGE_2_ actually leads to increased fungal germination and invasive hyphae (Fig 6B). Additional experiments also demonstrate that PGE_2_ injection does not increase survival of either wild-type or neutrophil-defective larvae not treated with indomethacin (Fig 6A, S7C Fig). These data suggest that the level of PGE_2_ during infection must be strictly controlled as too much PGE_2_ is also detrimental to control of *A. fumigatus*. Collectively, our findings demonstrate that COX-mediated PGE_2_ production and signaling via the EP2 receptor promotes phagocyte-mediated control of *A. fumigatus* germination and hyphal growth (Fig 7).

**Fig. 6.**
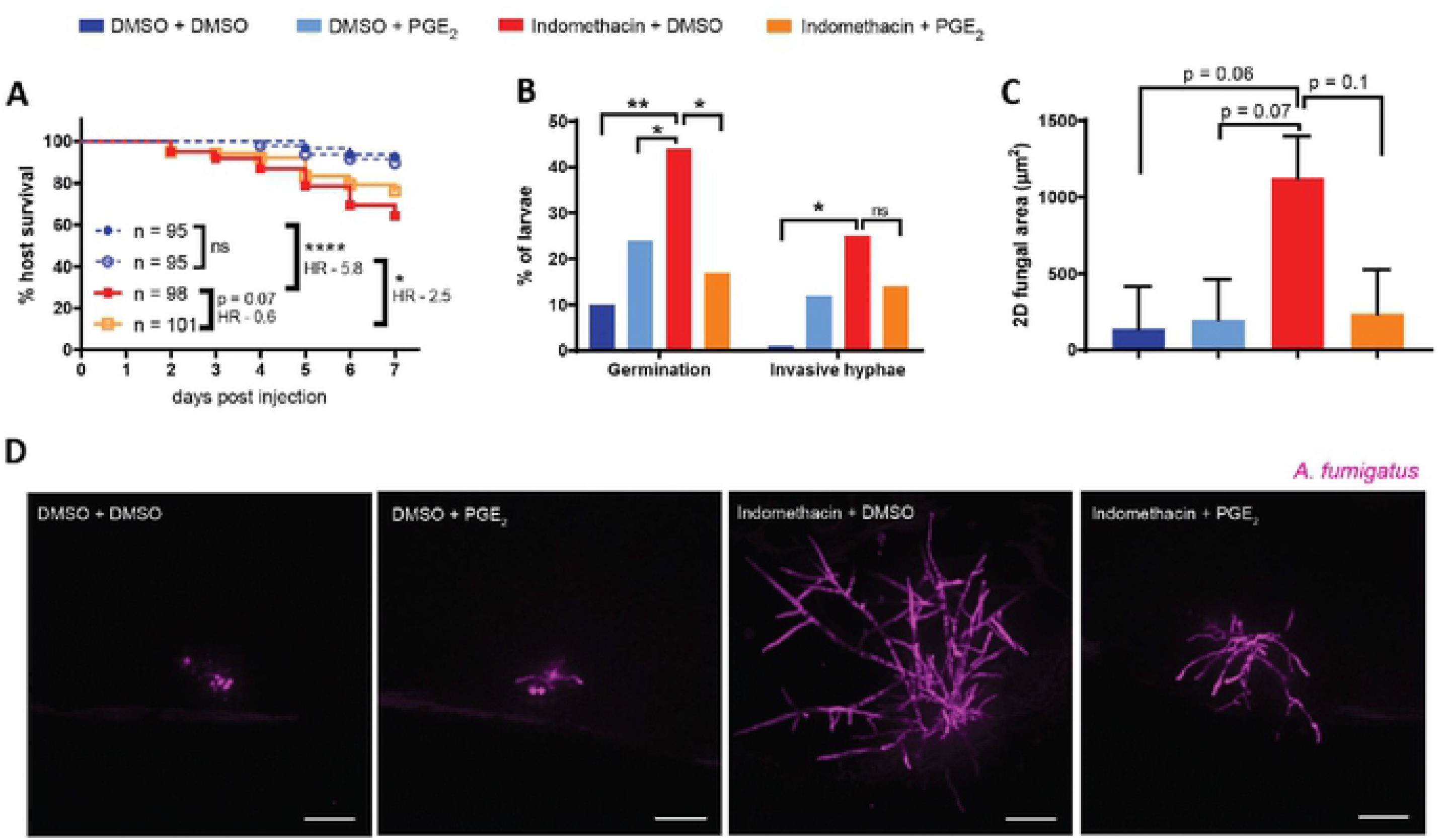
Exogenous PGE_2_ can rescue the indomethacin-mediated increase in fungal germination and hyphal growth. Larvae were injected with mCherry-expressing TBK5.1 (Af293) spores and exposed to 10 μM indomethacin or DMSO vehicle control at 2 dpf. At 1 dpi, larvae were injected with 10 μM PGE_2_ or DMSO vehicle control. **(A)** Survival of wild-type larvae was monitored. Cox proportional hazard regression analysis was used to calculate P values and hazard ratios (HR). Data are pooled from four independent replicates, and total larval N per condition is indicated in figure. Average injection CFUs: 40. **(B-D)** Larvae were imaged at 3 dpi. (B) Percentage of larvae with germination and invasive hyphae, and (C) 2D fungal area in each condition. Data pooled from three independent replicates, at least 8 larvae per condition, per replicate, P values calculated by ANOVA. (C) Bars represent pooled emmeans ± SEM. (D) Representative images showing hyphal growth in each condition. Scale bars = 50 μm.

**Fig. 7.**
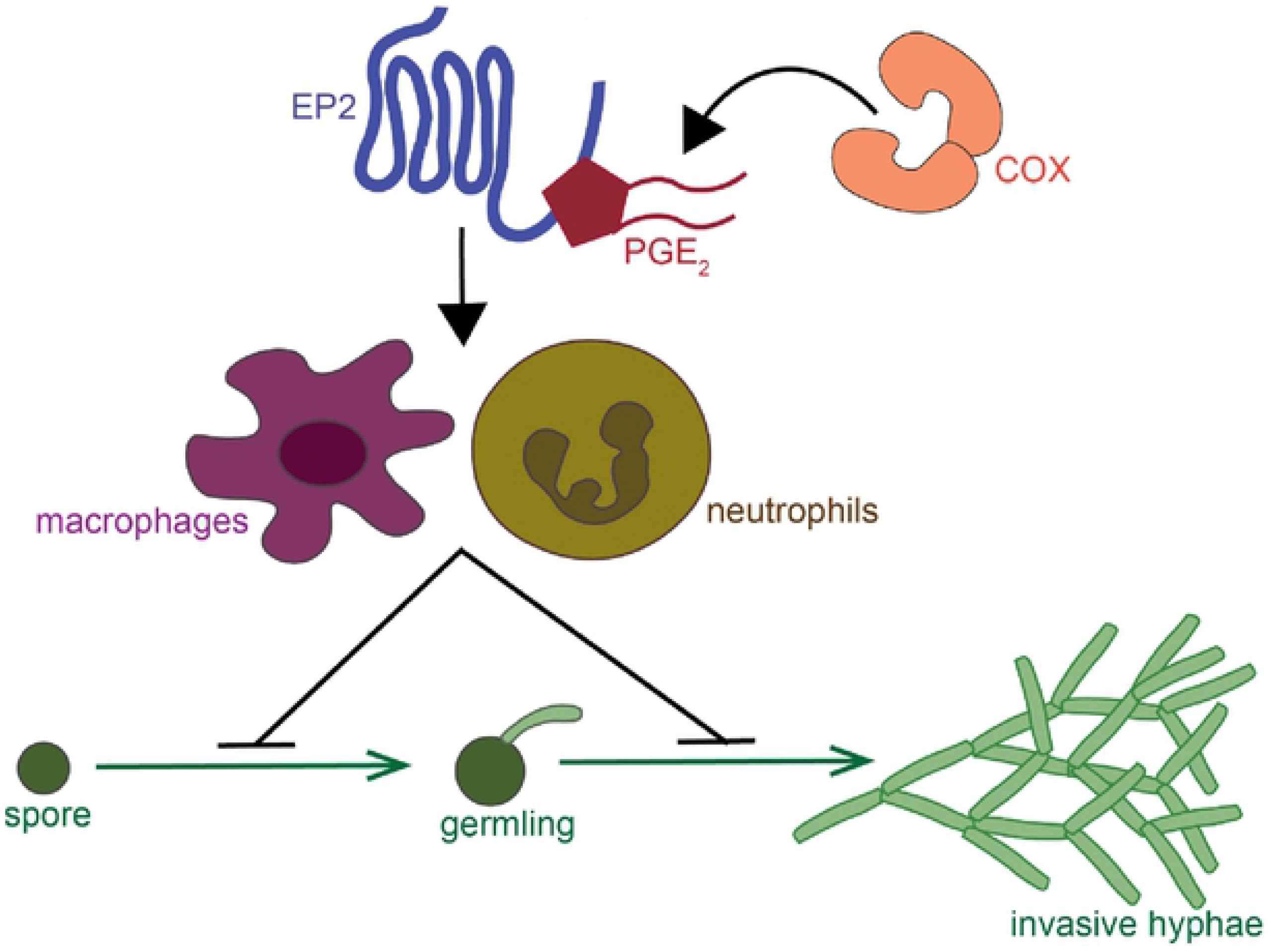
Model of cyclooxygenase signaling in response to *A. fumigatus* growth. COX enzymes produce prostaglandin signaling molecule, including PGE_2_. PGE_2_ can bind to the EP2 receptor to exert pro-inflammatory effects. Zebrafish larvae injected with *A. fumigatus*, this signaling activates both macrophages and neutrophils to inhibit spore germination and development of invasive hyphal growth.

## Discussion

Healthy immune systems can contain and kill *A. fumigatus* spores despite the fact that hundreds of spores can be inhaled per day. While the physiological role of macrophages and neutrophils in this context is well-appreciated, the molecular mechanisms that each of these cell types use to combat each stage of fungal pathogenesis are not fully understood. The critical step in *A. fumigatus* pathogenesis is the transition from dormant spore to hyphal growth, causing tissue destruction. Here, we used a zebrafish larva-*A. fumigatus* infection model to identify COX-PGE_2_ signaling as one mechanism that promotes control of this transition to invasive hyphae by both macrophages and neutrophils (Fig 7).

To investigate the role of COX signaling in phagocyte responses we used indomethacin, a pan-COX inhibitor, as well as COX1- and COX2-specific inhibitors. We find that activity of both enzymes promotes survival of *A. fumigatus*-infected larvae. Broadly, COX1 activity is involved in tissue homeostasis while COX2 is inducible and is involved in responses to inflammatory stimuli (6). However, evidence suggests that both isoforms are activated during inflammation. Mice lacking COX1 have impaired inflammatory responses (33), and COX1 is activated in response to LPS-induced inflammation in humans (34). Zebrafish have one functional isoform of COX1 and two functional orthologues of COX2: COX2a and COX2b (35). The role of COX1 in inflammatory responses in zebrafish is not fully understood, and the specificity of the chemical inhibitors for zebrafish COX2 isoforms are not know. Hence, we cannot conclude if phagocyte-mediated *A. fumigatus* control is via COX1, COX2a, COX2b, or a combination.

COX enzymes can produce prostaglandins in multiple cell types including epithelial cells, endothelial cells and fibroblasts, but infiltrating innate immune cells are the major source of these lipid signals during inflammation, including both macrophages (36) and neutrophils (11). In the current study, the source of prostaglandins is unknown. Both macrophages and neutrophils use COX signaling to combat *A. fumigatus*, however, we cannot rule out an additional role for other cell types in this signaling. One possibility is that PGE_2_ mediates crosstalk between these different cell types.

We also do not yet know what downstream effector mechanisms are activated in neutrophils and macrophages by COX signaling to target fungal growth. Prostaglandins can mediate endothelial cell permeability and facilitate immune cell infiltration, but we observe no difference in phagocyte recruitment when COX enzymes are inhibited (6, 37, 38). Prostaglandins can also regulate extracellular killing mechanisms in phagocytes such as reactive oxygen species (ROS) production, neutrophil degranulation, and extracellular trap (ET) formation (39–41). Further testing is required determine if these mechanisms are enhanced by COX signaling during infection with *A. fumigatus*.

COX enzymes catalyze the main regulatory step of prostaglandin synthesis: conversion of arachidonic acid to prostaglandin H_2_ (PGH_2_). PGH_2_ is then converted to one of the four major types of prostaglandins, prostaglandin E_2_ (PGE_2_), prostaglandin D_2_ (PGD_2_), prostaglandin I_2_ (PGI_2_) and prostaglandin F_2α_ (PGF_2α_) via different synthases. Perhaps the most studied prostaglandin is PGE_2_, due to its paradoxical immunomodulatory effects (42). PGE_2_ signals via four different receptors, EP1-EP4 which have different affinities for PGE_2_, and activate different downstream effects (26). PGE_2_ binds to EP2 with low affinity and generally evokes pro-inflammatory responses while it binds to EP4 with high affinity and activates anti-inflammatory responses (26). Therefore, the downstream effects of PGE_2_, whether pro- or anti-inflammatory, depend on local PGE_2_ concentration—which is controlled by its short half-life and the rate of synthesis by COX enzymes—and the cell type and EP receptor availability at the receiving end (42, 43).

We report here that PGE_2_ signaling, specifically through the EP2 receptor, promotes phagocyte control of *A. fumigatus* invasive hyphal growth. However, it was previously reported that PGE_2_ suppresses phagocytosis and microbial killing of fungal pathogens such as *P. brasiliensis* (12), *C. albicans* (13) and *C. neoformans* (14). These differences underline the idea that PGE_2_ can have both pro- or anti-inflammatory functions, can differentially impact diverse fungi, and that PGE_2_ levels must be tightly controlled during infection to promote infection clearance. Consistent with this idea, we report that while the effect of indomethacin on the control of fungal germination can be rescued by PGE_2_ supplementation, exogenous PGE_2_ in untreated infected larvae increases fungal growth. This could be due to elevated levels of PGE_2_ activating anti-inflammatory pathways and suppressing immune cell-mediated fungal growth control. In line with this possibility, PGE_2_ can drive resolution of inflammatory phenotypes through EP4 in zebrafish larvae during injury (28). Alternatively, the exogenous PGE_2_ might directly act upon *A. fumigatus* to promote germination. A previous study showed that exogenous PGE_2_ inhibited pigment formation in *A. fumigatus* hyphae which could affect invasive hyphal growth (24).

Fungal species can also synthesize their own lipid signaling molecules to modulate pathogenesis and immune responses (44, 45). For instance, *Cryptococcus neoformans* strains deficient in eicosanoid production have intracellular growth defects, which can be reversed by addition of exogenous PGE_2_ in a zebrafish larvae model (27). *A. fumigatus* PpoA and PpoC enzyme activity can produce prostaglandins and similar bioactive oxylipins that can affect *Aspergillus* virulence and development (24) and phagocytosis of conidia (46). However, we find that *A. fumigatus* Ppo enzymes do not affect fungal virulence in this larval zebrafish infection model, and that indomethacin treatment does not alter *A. fumigatus* spore germination *in vitro*.

Overall, PGE_2_ signaling must be well-orchestrated to elicit the desired anti- or pro-inflammatory effects. Hematopoietic stem cell transplant patients who are at high risk of developing IA harbor elevated levels of PGE_2_ (47), suggesting that modulating PGE_2_ or the downstream effects of PGE_2_ signaling may be a possible target for increasing control of these infections in patients. This study provides a first step towards understanding the function of this signaling in immune-mediated control of *A. fumigatus* infection.

## Materials and Methods

### Ethics Statement

Adult and larval zebrafish were maintained and handled according to protocols approved by the Clemson University Institutional Animal Care and Use Committee (AUP2018-070, AUP2019-012, and AUP2019-032). Buffered tricaine was used for anesthesia prior to any experimental manipulation of larvae. Adult zebrafish were euthanized with buffered tricaine and zebrafish embryos and larvae were euthanized at 4°C.

### Zebrafish lines and maintenance

Zebrafish adults were maintained at 28°C at 14/10 hr light/dark cycles. All mutant and transgenic fish lines used in this study are listed in Table 1 and were maintained in the AB background. Upon natural spawning, embryos were collected and maintained in E3 medium with methylene blue at 28°C. Embryos were manually dechorionated and anesthetized in 0.3 mg/mL buffered tricaine prior to any experimental manipulations. Larvae used for imaging were exposed to 200 μM N-phenylthiourea (PTU) starting at 24 hpf to inhibit pigment formation. Transgenic larvae were screened for fluorescence prior to experimentation. The *irf8* mutant line was maintained by outcrossing. *irf8*^+/−^ adults with fluorescent neutrophils (*Tg(mpx:mCherry)*) were in-crossed to generate *irf8*^+/+^, *irf8*^+/−^ and *irf8*^−/−^ larvae, and these larvae were screened for a high number of neutrophils to select *irf8*^−/−^ individuals (48). Genotypes were additionally confirmed at the end of the experiment where possible.

**Table 1.** Zebrafish lines used in this study.

| Zebrafish line | Reference |
| --- | --- |
| *irf8*<sup>-/-</sup> | (48) |
| *Tg(mpeg1:H2B-GFP)* | (49) |
| *Tg(mpeg1:H2B-mCherry)* | (50) |
| *Tg(mfap4:mTurquoise2)* | (51) |
| *Tg(lyz:BFP)* | (20) |
| *Tg(mpx:mCherry)* | (52) |
| *Tg(mpx:mCherry-2A-rac2D57N)* | (30) |

### Aspergillus fumigatus strains

Most experiments used Af293-derived strains TBK1.1 expressing YFP (19) or TBK5.1 expressing mCherry (20). Both of these strains behave like the parental Af293 strain in larval zebrafish (20). To test neutrophil-mediated killing, GFP-expressing TFYL49.1 (53) which was derived from the faster germinating CEA10 strain was used.

A Δ*ppoA*, Δ*ppoB*, Δ*ppoC* triple-mutant strain (∆*ppo*, TMN31.10) was used to test the role of fungal oxylipins. In these experiments the control comparison strain used was wild-type Af293. Briefly, a previously published Af293 ∆*ppoC pyrG1* strain TDWC3.4 (24) was used as the parental strain in which *ppoA* and *ppoB* were subsequently deleted, using the *A*. *parasiticus pyrG* marker and recyclable hygromycin resistance marker *hph* (54), respectively. All primers used for strain construction and confirmation are listed in S1 Table. DNA transformation constructs were created through double-joint PCR using published protocols (55). Protoplast generation and transformation were performed according to the previously published protocol (56). All transformants were first screened through PCR for incorporation of the construct and absence of the *ppo* gene. Southern blotting followed by hybridization of αP^32^-dCTP labeled 5′ and 3′ flank regions were used to confirm transformants with single integration (S2 Fig). The *ppoA* deletion construct was amplified from pDWC4.2 (GF ppoA del Cassette F and GF ppoA del Cassette R) and used to transform TDWC3.4, resulting in the prototroph Af293 Δ*ppoC* Δ*ppoA* TMN20. A deletion cassette for *ppoB* was constructed by fusing ~1 kb 5′ and 3′ flanking regions of the gene with the recyclable hygromycin B resistance gene *hph* from pSK529. *ppoB* deletion cassette was used to transform TMN20.11, resulting in the *ppo* triple deletion mutant TMN31. TMN31 transformants were subsequently grown on minimal medium with 0.1% xylose to recycle the hygromycin B marker.

### Spore preparation for injections

For injection preparation, 10^6^ spores were spread on solid glucose minimal media (GMM) 10 cm plates and grown at 37°C for 3-4 days. Spores were harvested by scraping using a disposable L-spreader and sterile water with 0.01% Tween. This spore suspension was passed through two layers of sterile miracloth into a 50 mL conical tube and topped to 50 mL. Spores were pelleted by centrifugation at 900 g for 10 min. The pellet was resuspended and washed in 50 mL of sterile PBS. The spores were again pelleted, resuspended in 5 mL of PBS, and filtered through another two layers of miracloth into a new conical tube. Spore concentration was enumerated using a hemocytometer. A final spore suspension of 1.5 × 10^8^/ mL was made in PBS and stored at 4°C for up to ~1 month.

### Live-dead spore labeling

Spores of *A. fumigatus* strain TBK1.1 were coated with AlexaFluor546 as described previously (20, 32). Briefly, isolated spores were incubated with biotin-XX, SSE (Molecular Probes) in the presence of 0.05 M NaHCO_3_ at 4°C for 2 hours. Spores were pelleted, washed first with 100 mM Tris-HCl at pH 8.0 to deactivate free-floating biotin and next with PBS, followed by incubation with streptavidin-AlexaFluor546 (Invitrogen). Spore concentration was enumerated and resuspended in PBS at 1.5 × 10^8^/ mL. Labeling was confirmed with fluorescence microscopy prior to injections.

### Zebrafish hindbrain microinjections

Larvae were injected with spores as described previously (16). Prepared spore suspensions at 1.5 × 10^8^/ mL were mixed at 2:1 with filter-sterilized 1% phenol red to achieve a final spore concentration of 1 × 10^8^/ mL. Injection plates were made with 2% agarose in E3 and coated with filter-sterilized 2% bovine serum albumin (BSA) prior to injections. Anesthetized 2 days post fertilization (dpf) larvae were placed on the agarose on their lateral side. A microinjection setup supplied with pressure injector, micromanipulator, micropipet holder, footswitch and back pressure unit (Applied Scientific Instrumentation) was used to inject 30-50 spores into the hindbrain ventricle of each larva. Larvae were injected with PBS as a mock-infection control. After injections, larvae were rinsed at least twice with E3 without methylene blue to remove tricaine and any free spores and were transferred to 96-well plates for survival and CFU experiments and to 48-well plates for imaging experiments.

### Clodronate liposome injections

*Tg(mpeg1:H2B-GFP)* larvae at 1.5 dpf were manually dechorionated and screened for GFP expression. 50 μL of clodronate or PBS liposomes (Liposoma) was mixed with 5 μL of filter-sterilized 1% phenol red and 2 nL was intravenously injected into the caudal vein plexus of GFP-positive larvae. After 24 hours, depletion of macrophages was confirmed by loss of GFP signal by screening with a fluorescent zoomscope (Zeiss SteREO Discovery.V12 PentaFluar with Achromat S 1.0x objective) prior to *A. fumigatus* infections.

### Morpholino injections

A *pu.1* (*spi1b*) morpholino oligonucleotide (MO) was previously published and validated (5’-GATATACTGATACTCCATTGGTGGT-3’) (ZFIN MO1-spi1b) (29) (GeneTools). Stock solutions were made by resuspension in water to 1 mM and kept at 4°C. For injections, the stock was diluted to 0.5 mM in water with 0.1% filter-sterilized phenol red and 0.5X CutSmart Buffer (New England Biolabs). A standard control MO (GeneTools) at 0.5 mM was used as an injection control. 3 nl of injection mix was injected into the yolk of 1-2 cell stage embryos. Efficacy of *pu.1* knockdown was determined by injecting MO into a macrophage-labeled zebrafish line and larvae were checked for fluorescence expression prior to *A. fumigatus* infections.

### Drug treatments

Infected larvae were exposed to the pan-COX inhibitor indomethacin (Sigma-Aldrich) at 10 μM, COX1 inhibitor SC560 (Cayman Chemical) at 5 μM, COX2 inhibitor meloxicam (Cayman Chemical) at 15 μM, EP2 receptor antagonist AH6809 (Cayman Chemical) at 5 μM or EP4 receptor antagonist AH23848 (Cayman Chemical) at 10 μM. These drugs were previously used in zebrafish larvae (21–23, 27, 28). The indomethacin concentration used was based on published results. For all other drugs, multiple concentrations were tested and the highest concentration of each drug that did not cause lethality or edema in uninfected larvae was used. 1000X stock solutions were made in DMSO and 0.1% DMSO was used as a vehicle control. After rinsing infected larvae, pre-mixed E3 with drug was added to dishes containing larvae. For survival and CFU assays, larvae were transferred to 96-well plates, one larva/well in 200 μL of drug/vehicle solution. For imaging experiments, larvae were transferred to 48-well plates with 500 μL of solution. All larvae were kept in the same drug solution for the entirety of the experiment unless otherwise noted.

### PGE_2_ rescue injections

A stock solution of 10 mM PGE_2_ (Cayman Chemical) was made in DMSO (23, 27). Prior to injection, the stock was diluted 100X in PBS and 1 μL was mixed with 9 μL of 1% phenol red for a final concentration of 10 μM. Wild-type larvae were injected with *A. fumigatus* and exposed to 10 μM indomethacin or DMSO vehicle control in 10 mL of E3 in 60 mm petri dishes. At 1 dpi, larvae were anesthetized and injected with 1 nL of 10 μM PGE_2_ or 0.1% DMSO into the hindbrain. After injections, indomethacin or DMSO control treatments were renewed, and larvae were transferred to 96-well plates for survival and 48-well plates for imaging experiments.

### CFU counts

Single larvae were placed in 1.7 mL microcentrifuge tubes in 90 μl PBS containing 1 mg/mL ampicillin and 0.5 mg/mL kanamycin, homogenized in a tissue lyser (Qiagen) at 1800 oscillations/min (30 Hz) for 6 min and spun down at 17000 g for 30 seconds. The whole suspension was spread on a GMM plate and incubated for 3 days at 37°C and the number of fungal colonies were counted. For all survival experiments, 8 larvae from each condition were plated immediately after injection to confirm actual injection dose and these numbers are reported in all figure legends. For CFU experiments to monitor fungal burden over time, 8 larvae were plated for each condition, each day and CFU counts were normalized to the average initial injection dose for that condition and graphed as percent initial spore burden.

### Live imaging

For daily imaging and PGE_2_ rescue experiments, individual larvae were removed from 48-well plates, anesthetized in tricaine and transferred to a zWEDGI device (57, 58). Larvae were imaged using a Zeiss Cell Observer Spinning Disk confocal microscope on a Axio Obsever 7 microscope stand with a confocal scanhead (Yokogawa CSU-X) and a Photometrics Evolve 512 EMCCD camera. A Plan-Apochromat 20x objective (0.8 NA) and ZEN software were used to acquire Z-stack images of the hindbrain area with 5 μm distance between slices. After imaging, larvae were rinsed with 200 μM PTU in E3 and put back in the same well (16). For AlexaFluor labeled live-dead staining, infected larvae at 2 dpi were mounted in 1% low-melting point agarose (Fisher BioReagents) in a 35mm glass-bottom dish (Greiner) and oriented laterally. Images were acquired using the same spinning disk confocal microscope with an EC Plan-Neofluar 40x objective (0.75 NA) with 2.5 μm distance between slices.

### Image analysis

All images were analyzed using Image J/Fiji (59). Presence of germination and invasive hyphae were manually analyzed. Any kind of hyphal growth, whether single or branched was considered an incidence of germination, while the presence of branched hyphae was considered an incidence of invasive hyphae. Maximum intensity projections were used to measure the 2D fungal area after thresholding the fluorescent intensity. The number of recruited phagocytes were counted across z-stacks using the Cell Counter plugin. For 2D phagocyte cluster area, the polygon selection tool was used to select and measure the area of macrophage cluster from maximum intensity projection of z-stacks. Images from the same experiments were used to quantify incidence of germination and invasive hyphae, fungal area, and phagocyte recruitment. To quantify live versus dead spores, images were blinded and processed with bilinear interpolation to increase pixel density two-fold. Cell Counter was used to manually count the number of live and dead spores in z-slices. All displayed images were processed in Fiji with bilinear interpolation to increase pixel density two-fold and are displayed as maximum intensity projections of z-stacks. Live versus dead spore images were additionally processed with a gaussian blur (radius = 1) to reduce noise.

### Aspergillus fumigatus in vitro germination assay

1 × 10^6^ TBK1.1 spores were inoculated into a flask containing 3 ml RPMI 1640 medium with HEPES (Gibco) containing 2% glucose in a 37°C shaker at 100 rpm in triplicate. Every 2 hours, 10 μL of the spore suspension was pipetted on to a microscope slide with a cover glass and imaged under the same Zeiss spinning disk confocal microscope using a Plan-Apochromat 20x objective (0.8 NA). At least 10 fields were captured for each sample at each time point and imaging was continued until 8 hours post seeding. These images were blinded prior to analysis and the number of germinated and non-germinated spores were counted using the Fiji Cell Counter plugin.

### Statistical analysis

For all experiments, pooled data from at least three independent replicates were generated and total pooled Ns are given in each figure. Statistical analyses were performed using R version 4.1.0 and graphs were generated using GraphPad Prism version 7 (GraphPad Software). Larval survival data and cumulative appearance of larvae with germination or invasive hyphae were analyzed by Cox proportional hazard regression. Calculated experimental P values and hazard ratios (HR) are displayed in each figure. HR defines how likely larvae in a particular condition will succumb to the infection compared to control larvae. Occasionally, the indomethacin drug lost efficacy and did not cause the previously observed and confirmed survival defect. Any such replicates were omitted from the final statistical analysis. CFU counts, spore killing, fungal area, phagocyte numbers and cluster area, and comparisons of day of onset of germination and invasive hyphae were analyzed with analysis of variance (ANOVA). For each condition, estimated marginal means (emmeans) and standard error (SEM) were calculated and pairwise comparisons were performed with Tukey’s adjustment. The graphs of number of phagocytes, phagocyte cluster area and 2D fungal area over the infection period show values from individual larvae over time as individual lines, and bars represent pooled emmeans ± SEM. Data points in dot plots represent individual larvae and are color-coded based on replicate and bars represent pooled emmeans ± SEM. For the *in vitro* germination assay, data were pooled from two independent replicates, each consisting of three technical replicates. Statistical analysis was performed by Student’s T-test with Holm-Sidak method using GraphPad Prism version 7, and data points represent pooled means ± SEM.

## Acknowledgments

We thank David Tobin for providing the *mfap4:mTurquoise2* zebrafish line. We thank Celia Shiau for providing the *irf8* mutant zebrafish line. We thank Anna Huttenlocher for sharing all other transgenic zebrafish lines. We thank members of the Rosowski Lab and Keller Lab for helpful discussions.

## Supporting Information

**S1. Fig. Both cyclooxygenase-1 and −2 signaling contribute to survival of infected larvae.** Larvae were injected with TBK1.1 (Af293) *A. fumigatus* spores at 2 dpf and were exposed to (A) 5 μM SC560, (B) 15 μM meloxicam, or DMSO vehicle control. Survival was monitored for 7 days. Cox proportional hazard regression analysis was used to calculate P values and hazard ratios (HR). Data are pooled from three independent experiments and the total larvae number is indicated in each figure. Average injection CFUs: (A) 35, (B) 30.

**S2. Fig. Southern blot analyses of strains created in this study.** Confirmation of (A) TMN20 Af293 Δ*ppoC* Δ*ppoA* double mutant and (B) TMN31 Af293 *ΔppoC* Δ*ppoA ΔppoB* triple mutant. Restriction enzyme digestion, southern blotting and hybridization were performed as mentioned in the Materials and Methods. Double and triple *ppo* mutants were created in sequence. Hybridization of αP^32^-dCTP labeled 5′ and 3′ flank regions were used to confirm transformants. The parental strain and the size of DNA fragments used to probe for southern blotting and hybridization are shown in each figure. P = parental strain; T = transformants.

**S3. Fig. Survival of phagocyte-deficient larvae after PBS mock-infection.** (A) Wild-type larvae injected with standard control MO or *pu.1* MO, (B) larvae injected with PBS liposomes or clodronate liposomes and (C) larvae with defective neutrophils (*mpx*:*rac2D57N)* or wild-type neutrophils were injected with PBS at 2 dpf. Survival was monitored in the presence of 10 μM indomethacin or DMSO vehicle control. Data are pooled from three independent experiments and Cox proportional hazard regression analysis was used to calculate P values and hazard ratios (HR).

**S4. Fig. Indomethacin does not affect *A. fumigatus* spore germination *in vitro*.** TBK1.1 (Af293) spores were inoculated into RPMI media in the presence of 10 μM indomethacin or DMSO vehicle control. Every 2 hours, an aliquot of spores was removed and scored for germination. Points represent pooled means ± SEM from two independent replicates and P values calculated by Student’s T-test.

**S5. Fig. Representative images of categories of *A. fumigatus* hyphal growth.** Wild-type larvae were injected with mCherry-expressing TBK5.1 (Af293) spores, exposed to 10 uM indomethacin or DMSO vehicle control and imaged at 1, 2, 3, and 5 dpi. Incidences of hyphal growth were scored a value of 1-4 depending on the extent of hyphae. Category 1: presence of one germ tube (white arrow). Category 2: presence of branched hyphae (open white arrow), yet small fungal bolus. Category 3: presence of spread-out invasive hyphae. Category 4: presence of severe invasive hyphae and tissue damage. Scale bars = 50 μm or 10 μm.

**S6. Fig. Indomethacin does not significantly impair neutrophil-mediated clearance of *A. fumigatus* hyphae.** Macrophage-deficient *irf8*^−/−^ or control (*irf8*^+/+^ or *irf8*^+/−^) larvae were injected with *A. fumigatus* TFYL49.1 (CEA10) strain and treated with 10 μM indomethacin or DMSO vehicle control. (A) Fungal burden was monitored by homogenizing individual larvae and quantifying CFUs at 1 and 2 dpi. CFUs from *irf8*^−/−^ were normalized to CFUs of *irf8*^+/+^/*irf8*^+/−^ at each dpi for each condition. Data were pooled from four independent replicates and P values calculated by ANOVA. (B) Larvae were monitored for survival. Data are pooled from five independent replicates and Cox proportional hazard regression analysis was used to calculate P values and hazard ratios (HR). Average injection CFUs: *irf8*^+/+^/*irf8*^+/−^ = 26, *irf8*^−/−^ = 20.

**S7. Fig. PGE_2_ injection does not alter phagocyte recruitment or rescue survival of neutrophil-defective larvae.** At 2 dpf, larvae were injected with TBK5.1 (Af293), followed by injection of 10 μM PGE_2_ or DMSO vehicle control at 3 dpf (1 dpi). The number of (A) macrophages and (B) neutrophils were enumerated at 3 dpi. Data are pooled from three independent replicates, at least 8 larvae per condition, per replicate. Each data point represents an individual larva, color-coded by replicate. Bars represent pooled emmeans ± SEM and P values were calculated by ANOVA. (C) Survival of injected and treated neutrophil-defective larvae (*mpx*:*rac2D57N*) was monitored. Cox proportional hazard regression analysis was used to calculate P values and hazard ratios (HR). Data are pooled from three replicates, at least 23 larvae per condition, per replicate. Average injection CFUs: 49.

**S1 Table. Primers used to construct *A. fumigatus Δppo* triple mutant strain.**

